# *v*LUME: 3D Virtual Reality for Single-molecule Localization Microscopy

**DOI:** 10.1101/2020.01.20.912733

**Authors:** Alexander Spark, Alexandre Kitching, Daniel Esteban-Ferrer, Anoushka Handa, Alexander R. Carr, Lisa-Maria Needham, Aleks Ponjavic, Mafalda Da Cunha Santos, James McColl, Christophe Leterrier, Simon J. Davis, Ricardo Henriques, Steven F. Lee

## Abstract

Super-Resolution (SR) Microscopy based on 3D Single-Molecule Localization Microscopy (SMLM) is now well established^1,2^ and its wide-spread adoption has led to the development of more than 36 software packages, dedicated to quantitative evaluation of the spatial and temporal detection of fluorophore photoswitching^3^. While the initial emphasis in the 3D SMLM field has clearly been on improving resolution and data quality, there is now a marked absence of 3D visualization approaches that enable the straightforward, high-fidelity exploration of this type of data. Inspired by the horological phosphorescence points that illuminate watch-faces in the dark, we present *v*LUME (Visualization of the Universe in a Micro Environment, pronounced ‘volume’) a free-for-academic-use immersive virtual reality-based (VR) visualization software package purposefully designed to render large 3D-SMLM data sets. *v*LUME enables robust visualization, segmentation and quantification of millions of fluorescence puncta from any 3D SMLM technique. *v*LUME has an intuitive user-interface and is compatible with all commercial VR hardware (Oculus Rift/Quest and HTC Vive, Supplementary Video 1). *v*LUME accelerates the analysis of highly complex 3D point-cloud data and the rapid identification of defects that are otherwise neglected in global quality metrics.

---

***v*LUME** is a point-cloud based 3D-SMLM data visualization tool able to render all pointillism-based multidimensional datasets. It differs from other 3D tools for 3D-SMLM visualization such as ViSP^4^ by providing a complete VR interactive environment and intuitive interface for life-scientists, dedicated to data visualization and segmentation. Users drag and drop multidimensional particle-list datasets into *v*LUME (.csv or .txt files, Fig. 1a), such as those generated by commonly used 3D-SMLM software^5,6^. This allows users to comprehend the spatial and temporal relation between points comprising a 3D structure. In time-lapse data, 3D reconstructions update for each time-point under user control. *v*LUME is based on the industry standard, crossplatform, Unity engine providing a high-performance rendering framework that updates and scales with future advances in graphics performance.

**Figure 1.**
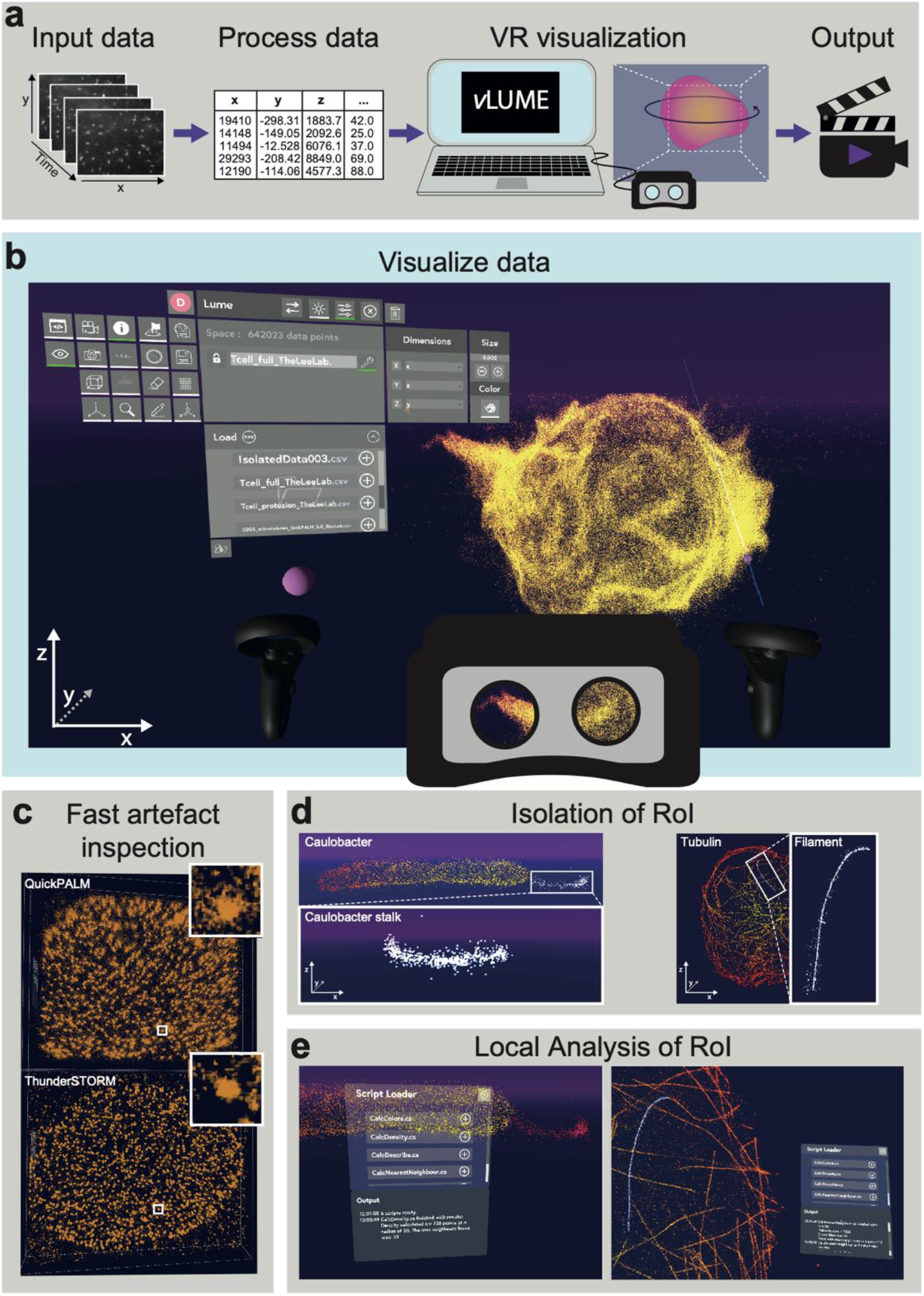
**a)** *v*LUME rapidly and simply takes large, multidimensional point-cloud datasets from 2D visualization into an immersive 3D VR environment through a systematic workflow: 1) multi-dimensional, SMLM image stacks are processed with any standard fitting algorithms providing multiparameter outputs as .csv or .txt files. 2) The resultant datasets can then be dragged and dropped directly into the *v*LUME software and instantly visualised in virtual reality. 3) By anchoring at user-defined waypoints around these data, a smoothly interpolated fly-through video can be created and exported providing the user with a tool to effectively communicate their discoveries. **b)** *v*LUME facilitates the 3D, VR visualization of millions of localizations, demonstrated by the super-resolved membrane of a T cell with *v*LUME. The accessible interface enables the user to customize their *v*LUME. The T cell is ~10 μm in diameter and has a isotropic resolution of ~24 nm. (FSA) **c)** Comparative inspection of artifacts introduced into data by localization fitting tools can be quickly performed. This can be seen by comparing localizations of Nuclear Pore Complexes fitted with both QuickPALM and ThunderSTORM algorithms. Scale the NPC of the ~100 nm in diameter, and the nuclear membrane ~20μm in diameter, data is taken over ~200 nm range axial range in z **d)** Selection and isolation of nanoscale, complex biological features can be easily achieved by the user. As examples, a stalk of *a Caulobacter crecentus* bacterium and a filament of a tubulin network were isolated and a Rol saved for further analysis. The microtubules tangle shows a region of ~ 20 μm × 30 μm (and about 500 nm in depth). The diameter section of a single microtubule is ~40 nm. **e)** Regions of interest can then be analyzed to instantly quantify desired properties using bespoke C# scripts (Ripley’s K, Local Density Plots, Nearest Neighbors and any others).

*v*LUME has 4 key features:

1. **Data Exploration and Comparison.** The configurable user interface allows researchers, without need for programming, to seamlessly switch back-and-forth from a global view of the entire captured sample, to detailed nanoscale views of molecular elements in any arbitrary orientation, faster than with conventional flat screens^7^. In doing so allows the easy local selection of data for further analysis (Supplementary Videos 2 and 3). The software can be used to leverage the human capacity to quickly interpret local features in these data, such as global and local artefacts (Fig. 1b) that are more difficult to trace by automated software without the ground truth being known^8^. In addition, it is easy to quickly evaluate and compare different processing software, side by side, *e.g*. QuickPALM *vs*. ThunderSTORM (Fig 1c). We include example data sets, with different sample types, using various SR methods, and from different international SR labs to demonstrate its broad applicability (Fig. 1b-e and Supplementary Information).
2. **Extracting 3D Regions of Interest (RoI) from complex data sets.** Complex biological interactions occur in intricate 3D geometries, with the evaluation of interaction data often requiring the extraction and analysis of specific sub-selections of a data set. To demonstrate this capacity of *v*LUME we carried out complex segmentation tasks where users need to identify and select small local features (tens to hundreds of localizations) in data of large dimensions (millions of localizations; Fig. 1d). A single microtubule can be easily extracted from a complex three-dimensional ‘web’ of microtubules within a eukaryotic cell (see also Fig. 1d). This process can be performed in less than one minute, and the RoI exported for further analysis (Supplementary Information). Once uploaded, these data subsets can be scaled, highlighted, coloured and selected in 3D via VR controls (Supplementary Video 4).
3. **Custom Analysis of user-defined subregions.** Quantitative bioimaging not only relies on high-quality images but quantitative evaluation using bespoke code. Recognizing this, we included a user-definable script interpreter written in the multi-paradigm language C# (Supplementary Information). These data can be easily evaluated to give the user instant quantitative feedback about the specific sub-region of their data set (Fig. 1e). We have included four widely-used analyses (Scripts in C#, Supplementary Information); Ripley’s K function^9,10^, nearest neighbour^11,12^, local density and the largest and shortest distances (these last both in Supplementary video 5).
4. **Exporting movies for publications and presentations.** As well as allowing customisation of data for presentation purposes *v*LUME also allows custom waypoints (user angle, pitch and yaw) to be defined simply in the VR environment (Fig. 1a) and to automatically generate a ‘fly-through’ video (Supplementary Video 6) to allow researchers to articulate their scientific discoveries.

In summary, *v*LUME provides a new immersive environment for exploring and analysing 3D-SMLM data. It enables imaging scientists with any level of expertise to make straightforward analytical sense of what is often highly complex 3D data. Full documentation and a link to the software (code availability) are included in supplementary information.

