## Supplementary Information for "*v*LUME: 3D Virtual Reality for Single-molecule Localization Microscopy"

Alexander Spark*, Alexandre Kitching*, Daniel Esteban-Ferrer*, Anoushka Handa*, Alexander R. Carr, Lisa-Maria Needham, Aleks Ponjavic, Mafalda Da Cunha Santos, James McColl, Christophe Leterrier, Simon J. Davis, Ricardo Henriques, Steven F. Lee.

### 1. SMLM Biological samples included with *v*LUME

We include 5 biological data sets, (11 files) from 4 different super-resolution microscopy laboratories obtained with a variety of different SMLM methods in the installation package of *v*LUME. The processed data sets (localization files) are saved in the comma separate values (.csv) file format and can be opened as point-clouds from the virtual environment of *v*LUME. They can be found in the \vLume_Data\StreamingAssets folder of the *v*LUME installation directory (see manual).

**T cell Plasma membrane (Lee Lab, Cambridge, UK)**

1.1 Jurkat T cell plasma membrane imaged in 3D using the Double Helix Point Spread Function (DH-PSF). The cell membrane was labelled using fluorescently tagged wheat germ agglutinin using PAINT, similarly to work previously described[1]. Each cell was imaged as series of 4 µm optical slices and reconstructed using overlapping fiducial alignment.[2] Localization data were processed using easy-DHPSF (10.1038/protex.2013.026). Fig. 1b shows the T cell within the virtual environment. The .csv file contains 3 columns; x position, y position and z position (/nm).
**Filename:** Tcell_full_TheLeeLab.csv

**Microtubules in U-2 OS cells (Ries Lab, Heidelberg, Germany)**

2.1 Microtubules in U-2 OS cells imaged with astigmatic dSTORM[3,4]. The microtubules were labelled with anti-alpha tubulin primary and Alexa Fluor 647-tagged secondary antibodies. The single emitters were processed using ThunderSTORM[3,5] (Fig. 1d and 1e, right). The comma delimited file contains 5 columns; frame number, x position, y position and z position (/nm) and intensity (/photons). These data are openly available (3).
**Filename:** U2OS_microtubules_ThunderSTORM_full_RiesLab.csv.

2.2 The same raw data of microtubules in U-2 OS cells as shown in 2.1 but processed using QuickPALM[3,6]. The comma delimited file contains 5 columns; frame number, x position, y position and z position (/nm) and intensity (/photons).
**Filename:** U2OS_microtubules_QuickPALM_full_RiesLab.csv.

2.3 Single microtubule obtained from a subset selection of sample 2.1 using *v*LUME (see Fig. 1d, right and Supplementary Video 2). The comma delimited file contains 5 columns; frame number, x position, y position and z position (/nm) and intensity (/photons).
**Filename:** U2OS_microtubules_ThunderSTORM_single_RiesLab.csv.

**Nuclear Pore Complex in U-2 OS cells (Ries Lab, Heidelberg, Germany)**

3.1 Nuclear Pore Complex (NPC) in U-2 OS cells imaged using astigmatic dSTORM[3,4]. The NPCs were labelled using Nup107-SNAP-BG tagged with AF647. The single emitters were processed using ThunderSTORM[3,5] (Fig. 1c, lower). The comma delimited file contains 5 columns; frame number, x position, y position and z position (/nm) and intensity (/photons). These data are openly available (3).
**Filename:** U2OS_NPC_ThunderSTORM_RiesLab.csv.

3.2 The same raw data of NPC in U-2 OS cells shown in 3.1 but processed using QuickPALM[3,6] (Fig. 1c, upper). The comma delimited file contains 5 columns; frame number, x position, y position and z position (/nm) and intensity (/photons). These data are openly available (3).
**Filename:** U2OS_NPC_QuickPALM_RiesLab.csv.

***Caulobacter crescentus* Bacteria (Moerner Lab, Stanford, USA)**

4.1 Colonies of the wild type (CB15N) strain of *Caulobacter crescentus* were grown overnight in a manner previously described[7] (Fig. 1d and 1e left). The bacteria were covalently labelled with N-hydroxysuccinimide functionalized rhodamine spirolactam via a coupling reaction with the exposed amines on the bacteria surface[7] and imaged in 3D using the DH-PSF and processed using bespoke software. The comma delimited file contains 3 columns; x position, y position and z position (/nm).
**Filename:** CaulobacterC_2_R09_MoernerLab.csv.

4.2 A single *Caulobacter crescentus* bacterium showing the stalk, imaged in 3D using the DH-PSF. The comma delimited file contains 3 columns; x position, y position and z position (/nm).
**Filename:** CaulobacterC_2_R23_MoernerLab.csv.

4.3 Stalk of a *Caulobacter crescentus* bacterium obtained using the data isolation tool in *v*LUME from dataset 4.1. The comma delimited file contains 3 columns; x position, y position and z position (/nm).
**Filename:** CaulobacterC_2_R09_stalk_MoernerLab.csv.

**Two-channel image of microtubules and clathrin in COS cells (Leterrier Lab, Marseille, France)**

Two channel image of microtubules and clathrin in COS cells imaged using astigmatism and with both DNA-PAINT and STORM labelling strategies[8]. Briefly, cells were labelled with rabbit anti-clathrin and mouse anti-tubulin antibodies, and secondary antibodies coupled to DNA-PAINT docking strands (Ultivue). The cell was imaged on a commercial N-STORM microscope (Nikon) in buffer containing imaging strands coupled to far-red (clathrin) and red (microtubules) fluorophores by alternating frames. Single emitters were detected using the N-STORM software (Nikon) and processed using ThunderSTORM.

5.1 The first channel is a COS cell labelled for microtubules imaged in 3D (Fig. S1, purple). The file contains 6 columns; frame number, x position, y position and z position (/nm), lateral uncertainty (/nm) and axial uncertainty(/nm).
**Filename:** COS_microtubules(ch1)_LeterrierLab.csv.

5.2 The second channel is labelled clathrin in COS cells imaged in 3D (Fig. S1, green). The file contains 6 columns; frame number, x position, y position and z position (/nm), lateral uncertainty (/nm) and axial uncertainty(/nm).
**Filename:** COS_clathrin(ch2)_LeterrierLab.csv.

### 2. Large datasets visualization

*v*LUME can easily render large point-cloud datasets in real time (see manual for further information). As an example, the T_cell_full_TheLeeLab.csv file has ~650,000 single-molecule localizations and opens up in ~15 seconds on a Gigabyte Aero 15 laptop, with an Intel i7 (8th gen.) CPU, an nVIDIA Geforce RTX 2080 GPU and 32 GB of RAM. It keeps a constant refresh rate of ~80-90 Hz while navigating the VR environment, which is the industry standard to avoid motion sickness, ~2 million localizations can be rendered without a significant drop in performance using the same system, this scales with the graphics processing power of the system.

### 3. Arbitrary selection and isolation of a region of interest (RoI) in large complicated datasets

Arbitrary selection and isolation of a local RoI is one of the tasks where VR provides an advantage over flat screens thanks to its intuitive navigation[9]. We performed the selection of a stalk in a *Caulobacter crescentus* bacterium and isolated its dataset (sample 4.3) (see Fig. 1d left and Supplementary Video 3) and a single microtubule from a complex tangle (sample 2.3) (see Fig. 1d right and Supplementary Video 2) in less than 1 minute (see manual). Using the selection feature also enables the application of bespoke C# scripts to perform analysis in any desired region.

### 4. Scripts in C# for local analysis

An important feature of *v*LUME the ability to perform local analysis on a subregion of point-cloud data (RoI). C# scripts can be programmed to perform any custom analysis once uploaded to the folder \vLume_Data\StreamingAssets\Scripts within the *v*LUME installation directory (.cs file extension). You can find further information on how program and use the scripts in the software manual.

We include four useful functions as .cs scripts for the *v*LUME interpreter that can be used for local cloud data-point analysis and that are often used in SMLM. You can apply them by selecting a RoI and toggling to the scripting menu (see manual). Note: depending on the complexity of the script you might see a busy indicator (out of the virtual environment) when called which is normal. Also be aware that some operations require more than the available memory particularly in large datasets (*i.e.* density plots with millions of points that need to compute every single distance from one point to another).

We include four scripts however it is our hope that we will nucleate communities to develop and share their own, included are:

**Ripley's K function (filename: CalcRipleysK.cs).** is a spatial point statistics analysis commonly used to evaluate clustering in SMLM data.[10,11] The script computes the Euclidian distance from a series of pairwise points in a RoI and counts the number of neighbours from a single point to a moving 3D radius (defined by the user and prompted as *RegionSize* and *RegionStep*). The input units need to be the same as the dataset. The output of the script is the L function, the linearized and localized Ripley's *K* function[11], for every *RegionStep* from 0 to *RegionSize*. The script also outputs a .txt file (named after the type of analysis and the time it was performed) in StreamingAssets\Scripts\Output saving the number of points in the RoI, the volume, region size and the step value. Then the computed, L(radius), and finally the position of every single point within the RoI.

Note: The function is only taken within the RoI as an isolated region and does not take into account any points in the periphery. Be aware that every single distance from point to point is computed, therefore large RoIs should be executed only in systems with large amounts of memory. The volume is computed as the smallest bounding box containing the whole dataset, which may be a suitable approximation for irregular volumes.

**Nearest Neighbour Plot (filename: CalcNearestNeighbour.cs).** A widely used analysis tool in SR[12,13]. The user is prompted for the value of the *ThresholdRadius* (that has to be inputted in the same units as the dataset) which will be used to compute the number of neighbours of every single point in the RoI up to the *ThresholdRadius*. The script then assigns a false colour depending on that number. These numbers are normalized with reference to the maximum number of neighbours, red being the lower density and blue the highest within a colour gradient (note: the colours will be plotted on top of the selection). The script also outputs a .txt file (named after the type of analysis and the time it was performed) in StreamingAssets\Scripts\Output, saving the number of points of the RoI, the radius tested, and the number of neighbours of every single point within the RoI together with its position.

As an example, the stalk of the *Caulobacter crescentus* bacterium (dataset 4.1) was analyzed (Fig. 1e, left and Fig. S2). We can clearly see that the number is larger in the core and lower in the surface as expected. We also analyzed the number of close neighbours of some of the NPCs (dataset 3.2) (Supplementary Video 5).

Note: The function is only taken within the RoI as an isolated region and does not take into account any points in the periphery. Be aware that every single distance from point to point is computed therefore, large RoIs should be executed only in systems with large amounts of memory.

**Calculate the Density of Points in RoI (filename: CalcDensity.cs).** We provide a very simple implementation that calculates the points within the RoI and divides this value by the volume of RoI to create the localization density. The result is printed within *v*LUME as points/volume using the units of the .csv dataset.

Note: The volume is computed as the smallest bounded box that can contain the selected dataset, this is only an approximation for irregular volumes.

**Calculate the Maximum and the Minimum Distances between points in RoI (filename: CalcShortAndFarDistances.cs).** For syntax purposes, we provide a very simple script that calculates the maximum and minimum distance between all the points within the RoI. The result is printed within *v*LUME using the units of the .csv dataset (Supplementary Video 5).

Note: Be aware that every single distance from point to point is computed therefore, large RoIs should be executed only in systems with large amounts of memory.

### 5. Exporting figures and videos

Videos (in .h264 format) and images (in .jpg format) can be captured from the *v*LUME environment and used for publication or presentation purposes. The fly-through video of the T cell (Supplementary Video 6), the images of the two channel Microtubule - Clathrin in COS cells (Fig. S1) and the Nearest Neighbour plot of the isolated *Calulobacter crescentus* (Fig. S2) were taken using these integrated features (*v*LUME manual). The .h264 videos can be opened and converted to other formats (*i.e.* .mov or .avi for larger compression ratios) in many video players (*i.e.* VLC).

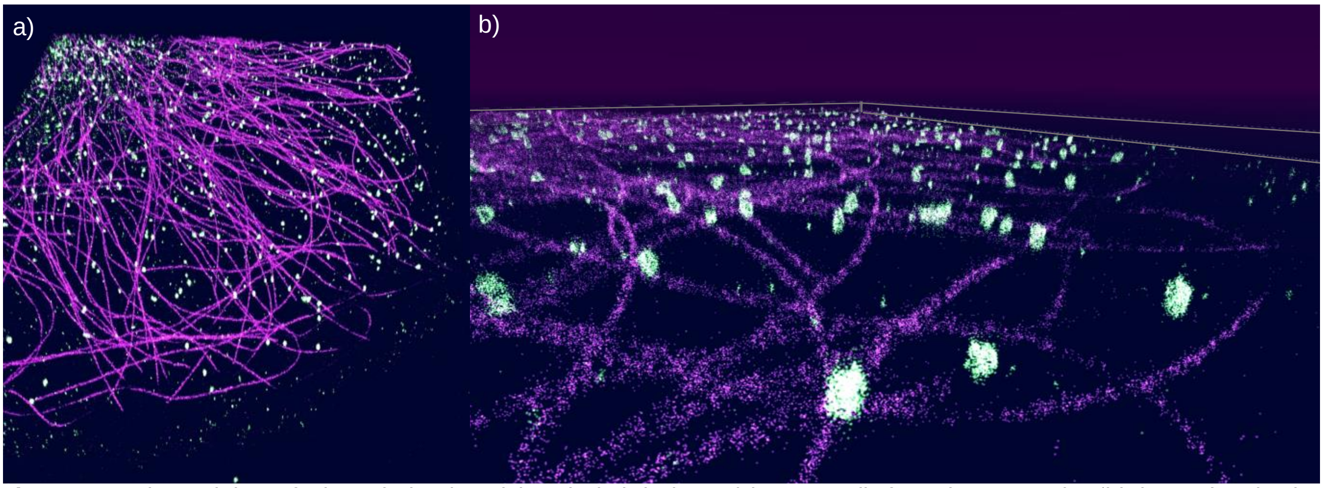


**Fig. S1** Two channel data of Microtubules (purple) and Clathrin (green) in COS cells from dataset 5. a) A 'birds- eye' projection of the two channels in *v*LUME. b) The same data set from a different point of view closer to the ground to show the 3D nature of the data. To achieve this superposition the first channel has to be opened in *v*LUME and the colour changed. Then the second channel also needs to be opened and changed in colour. Subsequently with simple data translation of one of the data sets the two axis need to be overlapped (this task is very simple in VR, see manual).

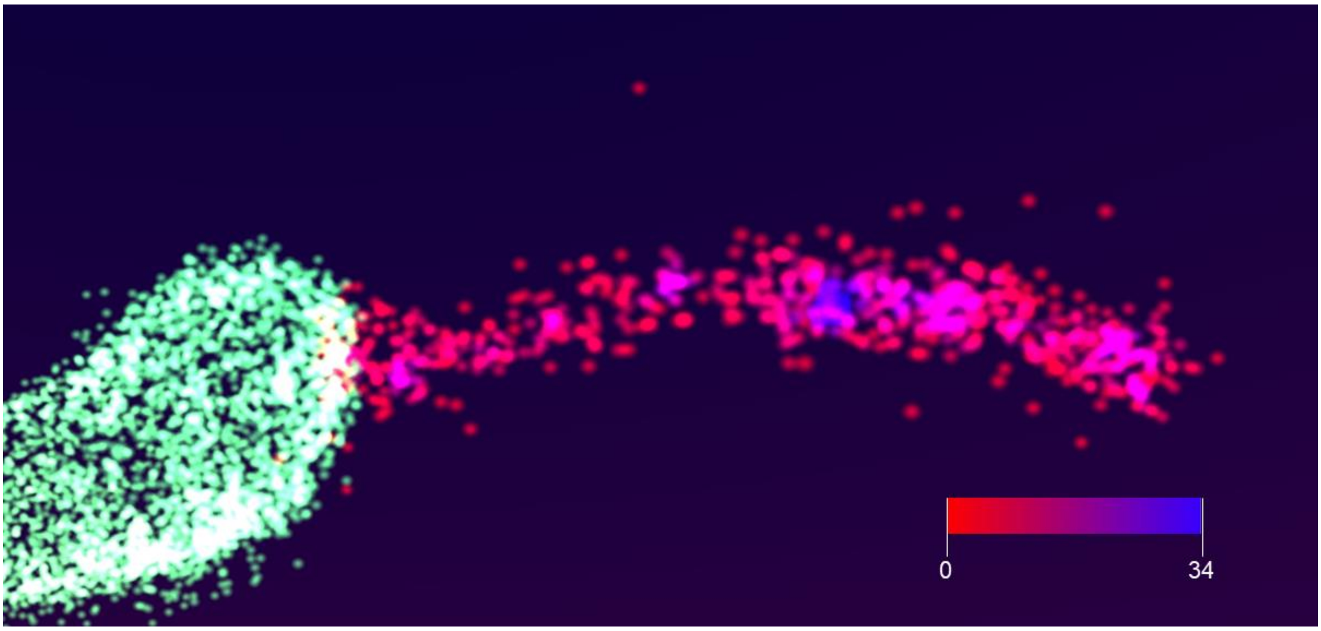


**Fig. S2** Nearest Neighbour plot using the C# script (CalcNearestNeighbour.cs) after selecting *Caulobacter crescentus'* stalk from dataset 4 of the Supplementary Information (see Fig. 1d and e, left). The red to blue gradient of the image shows an increasing density of nearest neighbours within a radius of 50 nm (user defined). The colour-gradient scale bar goes from 0 to 34 neighbours.

### 6. Supplementary Videos

**1. Overview of *v*LUME.** Overview of the main GUI and functionality in *v*LUME. The video shows a variety of data sets and the ease of going from the micron scale to molecular resolved regions (nanoscale).
**2. Selecting and annotating data.** Demonstration of a user isolating a single microtubule from a complex tangle in *v*LUME to be saved as isolated data.
**3. Loading and filtering data.** Demonstration of a user isolating a stalk from a pre-division stage of a *Caulobacter crescentus* in *v*LUME to be saved as isolated data.
**4. Manipulation of data.** Data manipulation features of *v*LUME on a single *Caulobacter crescentus*, showing the user actions simultaneously.
**5. Selecting Data and Running Scripts.** Maximum and Minimum distances and Nearest Neighbour script application to a NPC dataset.
**6. Outputting a video.** Setting waypoints in the 3D space using *v*LUME and saving these points as a video.

### 7. Acknowledgements

We would like to thank Dr. Marissa Lee, Dr, Joshua Yoon and Dr. Maurice Lee from the laboratory of W.E. Moerner (Stanford) for kindly providing the *Caulobacter crescentus* datasets 4 (Fig. 1d and e left). We would like to thank the laboratory of Jonas Ries (EMBL Heidelberg) for the publicly available Nuclear Pore Complex (NPC) dataset 3 shown in Fig. 1c and for the microtubule datasets 2 shown in Fig. 1d and e right. D. E-F would like to thank the European Union's Horizon 2020 research and innovation programme under the Marie Skłodowska-Curie grant agreement No 712949 (TECNIOspring PLUS) and the Agency for Business Competitiveness of the Government of Catalonia for the research funding leading to these results. We thank Royal Society for the University Research Fellowship of S.F.L., (UF120277) R.H funded by grants from the UK Biotechnology and Biological Sciences Research Council [BB/S507532/1], Wellcome Trust [203276/Z/16/Z] and core funding by the MRC Laboratory for Molecular Cell Biology, University College London [MC_UU12018/7].

### 8. Code availability

The updated versions of the software can be found at https://github.com/lumevr/vLume/releases. *v*LUME software for Windows (with manual, license, samples and scripts).

### 9. Competing interests.

While *v*LUME is free for academic use visualization software for SR microscopy, Alexandre Kitching and Alexander Spark are cofounders of Lume VR Limited; a company dedicated to creating human data interfacing software by leveraging the power of technology, such as immersive technologies. The remaining authors declare no competing interests.
